# A Statistical Model of Cell Wall Dynamics during Expansive Growth

**DOI:** 10.1101/346924

**Authors:** S. Lalitha Sridhar, J.K.E. Ortega, F.J. Vernerey

## Abstract

Expansive growth is a process by which walled cells found in plants, algae and fungi, use turgor pressure to mediate irreversible wall deformation and regulate their shape and volume. The molecular structure of the primary cell wall must therefore be able to perform multiple function simultaneously such as providing structural support by a combining elastic and irreversible deformation and facilitate the deposition of new material during growth. This is accomplished by a network of microfibrils and tethers composed of complex polysaccharides and proteins that are able to dynamically mediate the network topology via constant detachment and reattachment events. Global biophysical models such as those of Lockhart and Ortega have provided crucial macroscopic understanding of the expansive growth process, but they lack the connection to molecular processes that trigger network rearrangements in the wall. In this context, we propose a statistical approach that describes the population behavior of tethers that have elastic properties and the ability to break and re-form in time. Tether properties such as bond lifetimes and stiffness, are then shown to govern global cell wall mechanics such as creep and stress relaxation. The model predictions are compared with experiments of stress relaxation and turgor pressure step-up from existing literature, for the growing cells of incised pea (*Pisum sativus L.*), algae *Chara corallina* and the sporangiophores of the fungus, *Phycomyces blakesleeanus*. The molecular parameters are estimated from fits to experimental measurements combined with the information on the dimensionless number Π*_pe_* that is unique to each species. To our knowledge, this research is the first attempt to use a statistical approach to model the cell wall during expansive growth and we believe it will provide a better understanding of the cell wall dynamics at a molecular level.

## 1 INTRODUCTION

Algae, fungi, and plants have cell walls that act as an exoskeleton, providing shape, physical support, and protection from the external environment. Walled cells increase their volume and control their shape by a process called expansive growth. Expansive growth is central to development, morphogenesis, and sensory (growth and tropic) responses to environmental stimuli. During this process, the initial volume of the cell can increase by as much as 10,000 times. Interestingly, the cell wall (typically 0.1 – 1.0 µm in thickness) must perform two seemingly incompatible functions: (i) provide structural support and protection by employing strong and tough composites of complex polysaccharides and proteins and (ii) provide the ability to undergo extremely large plastic and elastic deformation under stress without rupturing. Plants, algae and fungi ^1,2^ all bring a similar solution to this dilemma; Water is first absorbed by osmosis through the plasma membrane to produce an internal turgor pressure (*P*) that stresses the wall. Biochemical reactions then alter the wall mechanical properties by breaking load-bearing bonds and causing a reduction in wall stress (stress relaxation). This further results in a drop in turgor pressure and a subsequently increases the water uptake that stretches the wall to increase the cell volume. Concurrently, new cell wall material is added to the inner wall to regulate its thickness. When water is not the limiting factor, the rate of expansive growth is thus directly related to the rate of wall stress relaxation ^3^.

Lockhart ^4^ derived the first global biophysical model and equations for expansive growth of walled cells, quantifying both the relative rates of water uptake and the coupled wall plastic deformation. In this model, the wall is seen as a so-called Bingham plastic fluid that can only flow (and permanently deform) once the turgor pressure (*P*) exceeds a critical value (*P_c_*). Lockhart’s equations have been used in many subsequent experimental investigations ^5–8^ that study the relationship between expansive growth rate, turgor pressure, and mechanical properties of the cell wall. They have also been used in their local form as constitutive equations to model complex growth situations^9^, including sporangiophore development ^10^ and “tip” growth morphogenesis ^11^. Though Lockhart’s equations for cell wall extension are attractive due to their simplicity, their applicability is limited because they do not consider elastic (reversible) wall deformations. Most importantly, Lockhart’s equation cannot describe the periodic stress and pressure relaxation that occur in cell walls ^1-3,7,12–16^ during expansive growth. In fact, elastic deformation is required to model wall stress relaxations ^13,17^ and the instantaneous changes in wall deformation after step changes in *P* ^18,19^. This issue was later addressed by the Ortega equations^15,16^ that describes the cell wall as a viscoelastic material ^17^ with a yield stress, commonly known as the Maxwell-Bingham viscoelastic model. In general, Ortega’s three equations (also known as the augmented growth equations) supplement Lockhart’s two equations with terms for transpiration and elastic deformation rates, and adds a third equation for the rate of change of the turgor pressure. Additionally, Ortega’s equations have been extended to include the pressure in the apoplasm (cell walls and xylem) of plant tissue and plant organs ^18^. Many experimental studies on plant cells in tissues, algal and fungal cells ^3,7,13,19–21^ have been successfully described and modeled with Ortega’s equations. Of particular interest is the fact that dimensional analysis of Ortega’s equations produces several dimensionless numbers, one of which describes the ratio of relative irreversible and reversible wall deformation rates, Π_pe_ ^22^. This number is central in categorizing different cell species in terms of their wall stress relaxation, elastic deformation and expansive growth ^23,24^. These biophysical models provide significant insight to understanding the mechanisms behind cell wall growth, but because they are macroscopically motivated, they do not provide information on the underlying molecular mechanisms. The growing (primary) wall is often envisioned as being composed of rod-like polysaccharide bundles of microfibrils, embedded in an amorphous network of polysaccharides and proteins. At the molecular level, the wall mechanical properties are regulated by facilitating the breaking of load-bearing bonds within the network, and making new bonds to incorporate new material into the wall that is released to the inner surface via exocytosis. Proteins such as expansin and enzymes ^25,26^ and changes in pH ^27,28^ have been shown to relax the wall stresses presumably by disrupting the load-bearing bonds. It is usually accepted ^29–32^ that the wall is able to support mechanical load via the tethers connecting microfibrils, while tether detachment is at the origin of wall extension. Experimental observations and results from plant, algal and fungal cells ^5,7,10,33,34^ have been qualitatively explained by this molecular mechanism and provides a molecular interpretation of the Lockhart and the Ortega equations. A comprehensive understanding of expansive growth however hinges on bridging the gap between the molecular mechanics and macroscopic behaviors. For instance, there is an abundance of literature on the molecular biology and biochemistry of plant and fungi cell walls during growth and sensory responses but macroscopic behaviors are described by continuum equations that do not have an immediate impact of the underlying molecular physics. A molecular scale version of Ortega’s equation would indeed allow researchers to bridge molecular effects, such as the effect of enzymatic or protein-driven growth regulation or environmental effects (light, gravity., etc) to macroscopic behavior and help explain intricate growth responses. For example, the presence of fibril slippage and reorientation sub-zones in the growth zone of the sporangiophores of *Phycomyces blakesleeanus* can qualitatively explain the observed helical growth behavior along the growth zone ^35,36^, but it has never been shown quantitatively. A molecular scale model can help clarify the associated mechanisms.

The objective of this paper is to address these needs by providing a route to *fundamentally understand how the organization and dynamics of transient tether-microfibril networks can lead to the emerging growth response of walled cells*. Here, we construct a theoretical framework that statistically describes the time evolution of a network based on molecular processes. The novelty of this work therefore lies in the ability to relate molecular properties, such as tether stiffness and bond lifetimes, to global wall mechanics, such as growth and stress relaxation, by a statistical description of the population behavior. The article is organized as follows. First the dynamics of molecular tethers and their mechanical properties are described, followed by a continuum statistical description of the tether population dynamics. This section is then followed by the application of the statistical model to describe global dynamics of the cell wall during steady growth, stress relaxation, and extension-growth response to a sudden increase (step-up) in turgor pressure. Previously published experiments of stress relaxation and extension-growth after pressure steps obtained from growing cells of incised pea stems (*Pisum sativus L.*), sporangiophores of the fungus *P. blakesleeanus,* and algal internode cells of *Chara corallina* are compared with model predictions and the importance of Π_pe_ is highlighted. Finally, the roles of the molecular parameters of the cell wall are discussed in the context of growth regulation that can offer crucial insights and potential grounds for future work.

## 2 A STATISTICAL MODEL OF THE CELL WALL

### 2.1. Cell Wall Geometry and Internal Architecture

The expansive growth of a walled cell is essentially driven by turgor pressure that causes the wall to expand (deform) in response to tensile stresses. The nature of these wall stresses and deformations however depends on the geometry of the cell, as well as the molecular, anisotropic structure of its wall and biochemical reactions that modify the mechanical properties of the wall. It is therefore not surprising that by combining these elements, plant and fungal cells can display a large variety of growth and morphogenetic events (twist, bend, differential growth) during their life time. We are interested in the way by which these processes occur at the molecular level and therefore need a more accurate description of the cell wall. We consider the primary cell wall to be an anisotropic network of unidirectional and long microfibrils that are interconnected by smaller, load bearing tethers as shown in Fig. 1. As the network is subjected to tensile stresses, the stiff fibrils undergo very little deformation, while tethers, due to their size and elasticity, are responsible for most of the wall’s elastic deformation. But the role of tethers is not limited to elastic deformations; their dynamics are at the source of irreversible deformation and growth. The molecular mechanisms of wall extensions can be generally understood as follows. As the wall is subjected to stress (a) tethers first undergo an elastic stretch which is at the origin of the wall’s elasticity, (b) if given enough time, these tethers may subsequently detach from a fibril, leading to intermittent events of stress relaxation in the wall and (c) detached tethers may finally re-attach to new microfibrils in a force-free configuration (see Fig. 1). This last event enables the network to preserve its mechanical integrity over time. The cell wall may therefore be seen as a highly dynamic network of tethers that continuously changes its topology during deformation.

**Figure 1:**
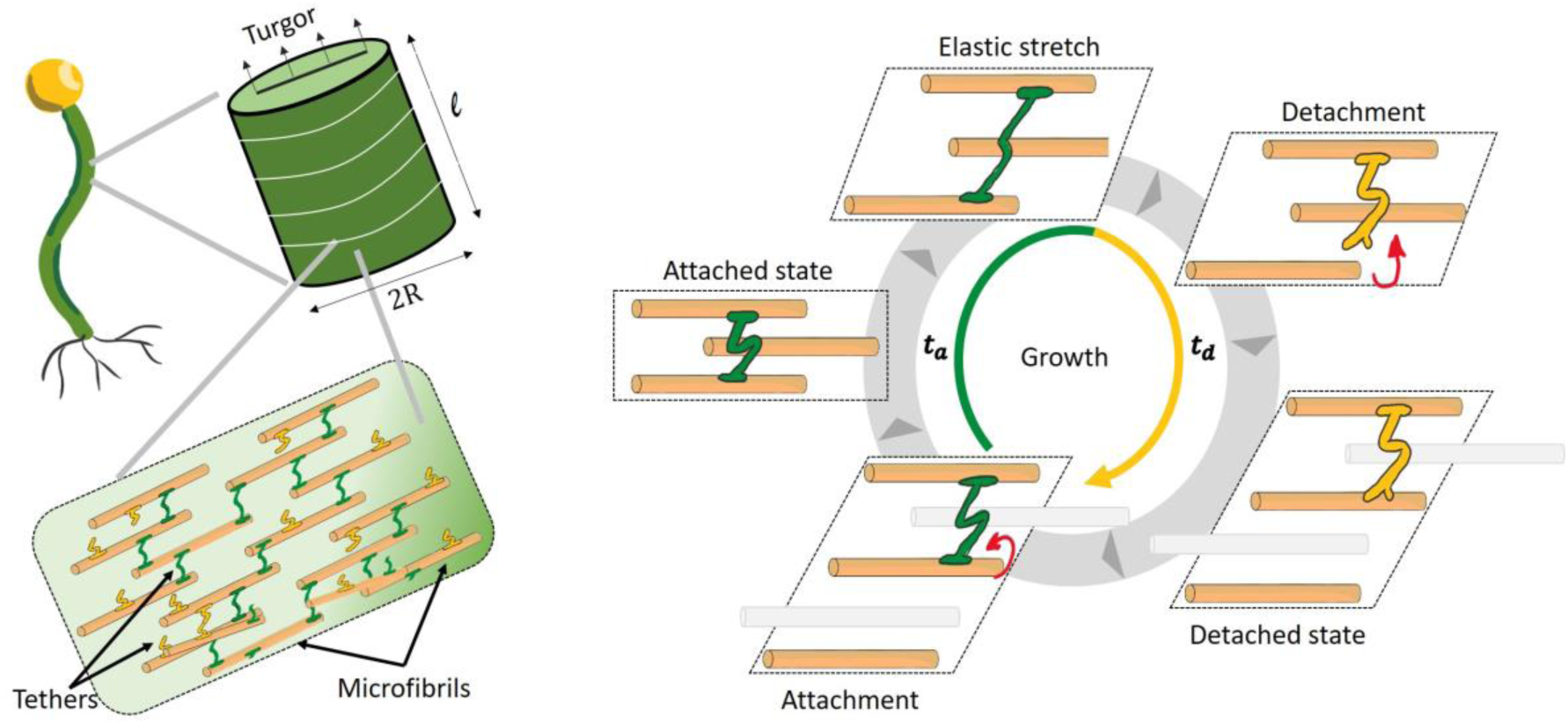
Left: Schematic of the expanding cylindrical cell wall composed of a dynamic network of microfibrils and tethers. Right: Illustration of the expansive growth process of the wall governed by elastic stretching of tethers in the attached state (green) with lifetime, t_a_ and relaxation due to detachment (yellow) that lasts for the lifetime, t_d_.

To focus on this molecular mechanism, we restrict our study to longitudinal growth of the cell wall. It is envisioned that the longitudinal growth occurs in the absence of radial expansion of the cylindrical wall structure. Thus, the total tensile force in the longitudinal direction of the wall is obtained from balance of forces on the cross-sectional surface of the cylinder as *F* = *P πR*^2^, where *P* is the turgor pressure and *R* is the cell radius. Therefore, it is sufficient to model a one-dimensional network of tethers connecting transversely oriented microfibrils subjected to the force, *F* (Fig. 1). We note here that the orientation of microfibrils need not remain in the transverse as shown in Fig. 1 and have in fact been postulated to reorient to produce helical growth in some cells ^35,36^. However, for the sake of simplicity we reserve such morphogenetic behavior of the cell wall for future investigations. In order to prevent the wall from thinning and rupturing during longitudinal expansion, new fibrils and tethers are deposited by the cell, as shown in Fig. 1 (grey microfibrils). Thus, the densities of microfibrils and tethers in the surface area of the primary cell wall are assumed to remain approximately constant during growth.

### 2.2. A Probabilistic View of Cell Wall Mechanics

Based on the above mechanisms, it is possible to derive a simple mathematical model of tether dynamics in the context of Poisson processes. For this, we consider that a tether can be found in two distinct states: attached and detached. The average lifetime of a tether in its attached and detached state can be described by the variables *τ_a_* and *τ_d_* respectively. Because the events of detachment and attachment are random in time, they constitute a Poisson process. The average values of the lifetimes can be related to average rates of occurrence of each event: the detachment rate, *k_d_* = 1/〈*t_a_*〉, and attachment rate, *k_a_* = 1/〈*t_d_*〉, where 〈. 〉 denotes average. Between times *t* and *t* + *δt*, the probability of detachment of an attached tether is given by *k_d_δt*, where *δt* is an infinitesimal time interval. Similarly, the probability that a detached tether re-attaches to the fibril in this time span is *k_a_δt*. It is then possible to calculate the probabilities of survival in the attached (*P_a_*) and detached (*P_d_*) states that decay with time as ^37^

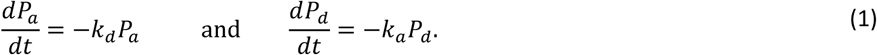

In other words, the longer a tether remains in a particular state, the more likely that it will change states. As permanent deformation of the cell wall occurs only after the wall stress is above the yield stress ^4^, we introduce a critical force, *f_c_*, on each tether below which detachment does not occur. Above the critical force, however, the tethers detach at an average rate *k_d_* such that:

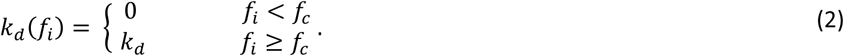

We will see that the above condition necessitates a net yield force on the wall that overcomes the critical force of all attached tethers in order to achieve permanent deformation.

Let us now consider a population of n tethers under the effect of a macroscopic force, *F*. In this case, it is possible to define the physical state of a tether (indexed by the integer *i* ∊ [1, *n*]) by a state variable *s_i_* that takes the value of 1 when it is attached and the value of 0 when detached. It is therefore possible to construct a simple computational algorithm that can predict the evolution of this tether population in time by integrating the state variables *s_i_*(*t*) over time. To understand how this process affects the wall mechanics, consider now that the tethers are represented by linear springs of stiffness *K* that are, when attached, subjected to an elongation *δ_i_*, with respect to their rest configuration. The elastic force in tether *i* is therefore given by Hooke’s law in the form *f_i_* = *Kδ_i_* and the condition of mechanical equilibrium implies that the macroscopic force is related to tether forces by:

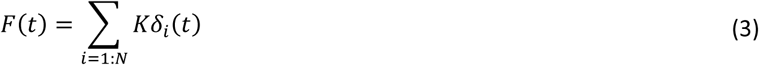

where we implicitly assumed that tethers act as a parallel assembly of springs. We will see below that if a tether is detached, its elongation vanishes, which means that it does not participate in resisting the force F. If the microfibrils are now separated at a macroscopic velocity *v*(*t*), the tether elongation is given by

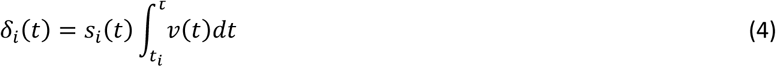

where *t_i_* is the time at which tether *i* attached to a microfibril and the presence of *s_i_* enforces that the stretch vanishes for a detached tether. When a stretched tether detaches, Eq. 3 therefore predicts a drop in macroscopic force of magnitude *Kδ_i_*(*t*). Thus, as tether detachment contributes to stress relaxation, the reattachment of new tethers causes the network to keep its connectivity and an effectively transfer the load between units. We note that tethers detach in a stretched state and re-attach in a stress-free state, i.e. *δ_i_*(*t_i_*) = 0. This “asymmetry” in force during the detachment and attachment events is responsible for permanent microfibril movement and wall deformation. Figure 2 provides an illustration of the mechanisms at play as the wall undergoes a deformation at rate *v*. We see that random tether attachment and detachment events results in changing the stretch distribution (represented by histograms) of the tethers over time. This distribution has a direct effect on the force *F* in the network according to Eq. 3. A Monte-Carlo simulation approach was constructed based on the above equations. The algorithm described in Appendix A.1 relies on a random sampling method and the probability criteria in Eq. 1. Numerical results based on this approach are given in the remainder of this study.

**Figure 2:**
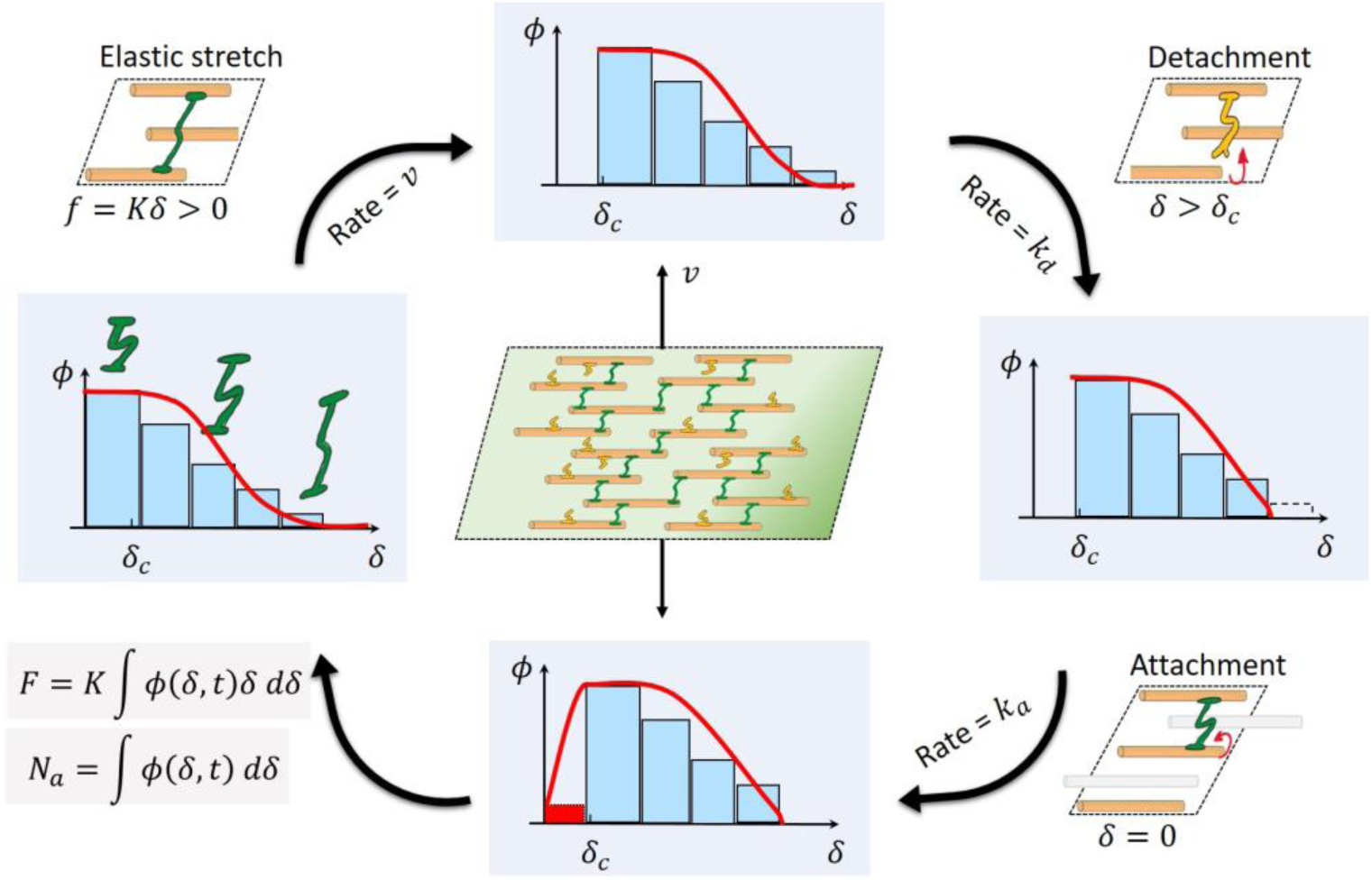
Illustration of stretch, detachment and attachment of tethers and the change in the tether population behavior depicted with histograms and approximated with the continuous distribution function ϕ(δ, t) when the size of the population is large. The total number of attached tethers N_a_ and wall force F are obtained from ϕ and its evolution.

### 2.3. Statistical Mechanics of Large Tether Populations

The probabilistic approach provides a comprehensive tool to study how a finite number of tethers and their behaviors affect a small portion of the wall mechanics. However, when the population becomes large, this approach becomes excessively costly, while at the same time, seem to converge towards more predictable and continuous trends. The mechanical response of large networks can instead be described by continuum mechanics, which may eventually take the form of macroscopic models such as the Ortega equations. We here propose to build a bridge between the scale of a few tethers to the full population using statistical mechanics, for which a framework in similar transient networks was previously discussed ^38,39^.

The transition between discrete and continuum is accomplished via the introduction of a stretch distribution function *ϕ*(*δ*, *t*) where *δ* is a continuous random variable representing the tether elongation. The distribution function is so defined that, the quantity, *ϕ*(*δ*, *t*)*dδ*, denotes the number of attached tethers with an elongation value in the interval *δ* and *δ* + *dδ*, at any given time *t*. Its integral over all elongations *δ* (denoted here as the configuration space) then gives the total number of attached tethers, *N_a_*(*t*). The total force, *F*(*t*), due to the tethers on the fibrils is now given by an integral form of Eq. 3. These conditions are expressed mathematically by

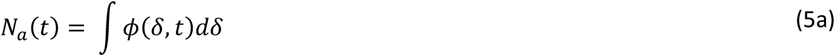

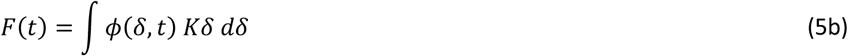

As the tether population is dynamic, the function *ϕ*(*δ*, *t*) evolves in time due to: (a) the macroscopic deformation of the network that causes attached tethers to elongate at rate 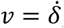, and (b) the random detachment and attachment of tethers at rates *k_d_* and *k_a_*, respectively. Let us decompose the rate of change of *ϕ*(*δ*, *t*) into

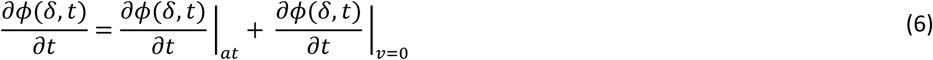

The first term in Eq. 6 denotes the evolution in *ϕ*(*δ*, *t*) due to a purely elastic wall deformation when the population of attached tethers (subscript *at*) remains unchanged, i.e. there are no attachments and detachments (*k_a_* = *k_d_* = 0). The second term denotes the change in tether distribution due to tether kinetics when the wall is not growing, i.e. the microfibrils are fixed in position and *v* = 0.

To compute the first term, let us consider a population of attached tethers in an arbitrary range of tether elongation, Ω*, that neither attach or detach. Due to macroscopic deformation of the network, all attached tethers in this population are stretched, thereby shifting the elongation range, Ω* (see illustration in Fig. 2). However, if we follow the state of this tether population during deformation, the number of attached tethers remains constant in time as there are no attachments or detachments. Mathematically, this can be described using the “material” time derivative (denoted by *D* (.)/*Dt*) as

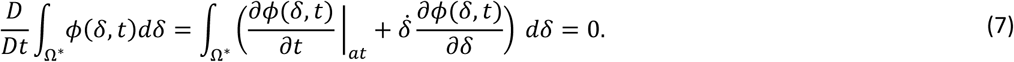

where the integration of *ϕ*(*δ*, *t*) in the range, Ω*, gives the number of attached tethers in that range. As the above equation holds true for any arbitrary range, Ω*, it can be localized to give

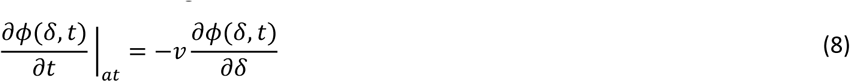

To compute the second evolution term of Eq. 6 at *v* = 0, we first assume that the effective rate of detachment of tethers with elongation, *δ*, is proportional to *ϕ*(*δ*, *t*). Furthermore, we assume that re-attachment of detached tethers occurs at a stress-free configuration that is described by the probability density function, *p*_0_(*δ*). The evolution equation is thus given by

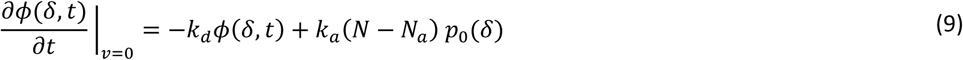

where N is the total number of tethers in the network and *N* − *N_a_* is the number of detached tethers at any time *t*. Combining Eqs. 6,8 and 9, we get the evolution of *ϕ*(*δ*, *t*) as

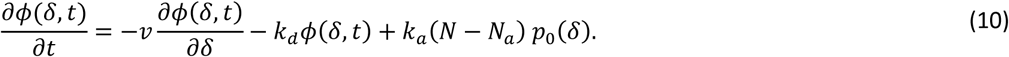

Thus, the Eqs. 5 and 10 provide the general mathematical framework needed to relate the tether population behavior to the macroscopic wall force, *F*(*t*), and growth rate, *v*(*t*).

In this paper, we assume for simplicity that the rates *k_a_* and *k_d_* are constants in time. While *k_a_* is also independent of tether elongation, *k_d_* follows the rule described in Eq. 2 for a critical tether elongation, *δ_c_* = *f_c_*/*K*. We also assume that the probability density, *p*_0_(*δ*), is a Dirac delta function centered at *δ* = 0, so that attachment always occurs at a stress-free state. By calculating the time rate of total network force, 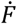, using Eqs. 5 and 10, we can obtain a macroscopic evolution equation (see Appendix B.1) given by

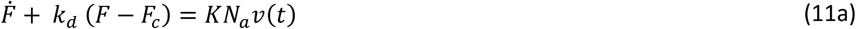

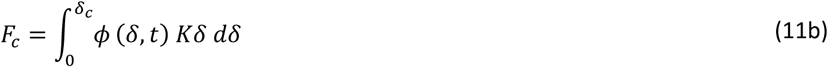

where *F_c_* is the overall force from the “static” (non-detaching) tether population with elongation below *δ_c_*. The creep velocity *v*(*t*) is equivalent to the longitudinal growth rate of the cell wall driven by the wall force *F* and is governed by the molecular parameters, *k_d_*, *k_a_*, *K*, and *f_c_*.

To illustrate the statistical mechanics model with the discrete and continuum approaches, let us consider the case when a constant force, *F*, is applied to the network. If the force is large enough to induce a tether force *f* that is larger than its critical value *f_c_*, permanent deformation ensues as a result of tether detachment. For given values of the molecular parameters *k_d_*, *k_a_*, *K*, and *f_c_*, (see Fig. 3 caption) Monte Carlo simulations predict an approximately constant rate of change of separation distance between microfibrils, i.e. wall length. The fluctuations in the wall length arise due to discrete events of tether detachment and attachment in a population size of *N* = 100. The fluctuations are observed to reduce with increase in population size to *N* = 1000 and *N* = 10,000. The histograms shown in Fig. 3 correspond to the tether elongation of the attached population after it reaches an approximately steady state in time. This steady state, however, is an outcome of a dynamic equilibrium of deformation, detachment and re-attachment of tethers. In the case of a large population size where the continuum assumption is justified, steady state implies that the distribution function, *ϕ*(*δ*, *t*) remains constant in time, i.e. *∂ϕ*/*∂t* = 0. The evolution Eq. 10 can then be solved analytically (see Appendix B.2) to give:

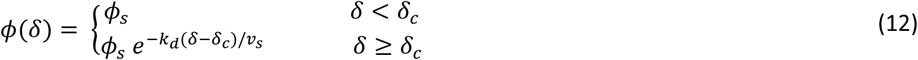

where *ϕ_s_* = *k_a_*(*N* − *N_a_*)/*v_g_* is constant in time. The steady state growth rate or creep velocity *v_g_*, is given by Eq. 11a, where 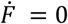 as the wall force is constant. The estimation of the wall deformation (irreversible) and the population distribution of tether elongation from the continuum approach are smooth and good approximations of large tether populations as demonstrated in Fig. 3.

**Figure 3:**
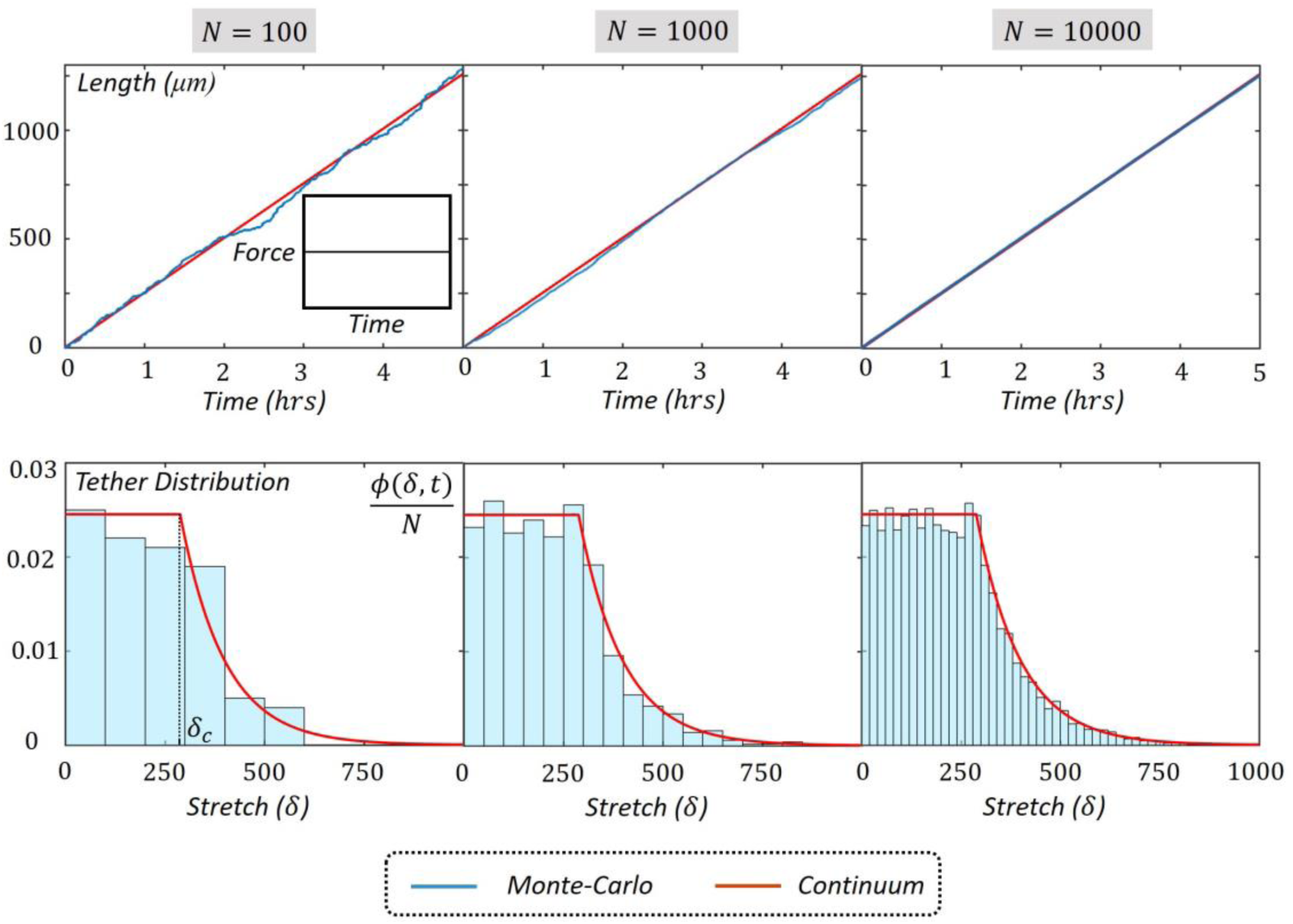
Comparison of predictions from discrete Monte Carlo simulation and the continuum models for increasing sizes of tether population, N = 100,1000 and 10000. Top: Growth of network length in time for a constant force, F = 1000 μN. Bottom: Distribution function of tether elongations, ϕ(δ, t), normalized to the population size, N at steady-state. The molecular parameters used for all simulations are k_a_ = 10 h^−1^, k_d_ = 2 h^−1^, NK = 5 Nm^−1^ and Nf_c_ = 1500 μN.

## 3 INVESTIGATING GLOBAL WALL MECHANICS

Interestingly, Eq. 11a has a very similar form as Ortega’s growth equation (AGE) that describes the rate of change of the cell volume as a function of turgor pressure, *P*. For a cell wall with negligible radial expansion, the cross-sectional area of the cell, *A*_0_ = *πR*^2^, remains constant and Ortega’s equation can be written as

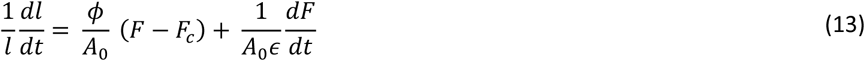

where *l* is the length of the cell wall, *ϵ* is the volumetric elastic modulus and the wall extensibility *ϕ* governs irreversible deformation. Comparing Eqs. 11a and 13, we find that the bulk wall properties in the Ortega equation are related to the molecular parameters as *ϵ* = (*KN_a_l*/*A*_0_) and *ϕ* = (*k_d_A*_0_/*KN_a_l*). Therefore, rewriting Eq. 11a to have a similar form as Eq. 13, we obtain the molecular version of the Ortega equation as

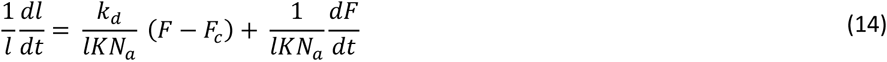

The stress relaxation time for the Ortega model (Maxwell-Bingham fluid)^17^ can be obtained as *t_R_* = 1/(*ϵϕ*) = 1/*k_d_*, while the average relative growth rate of the wall is given by *v*_s_ = *v*/*l*. The viscoelastic nature of the cell wall may then be characterized by the dimensionless Π_pe_ number ^23,24^ defined as

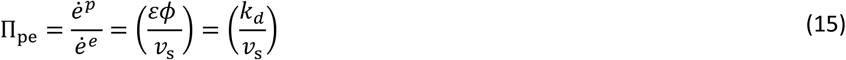

that describes the ratio of the relative plastic 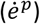 and elastic 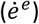 deformation rates of a viscoelastic cell wall. Recent studies have shown that this number may be a crucial descriptor of the chemo-rheology of the cell wall during expansive growth. The magnitude of Π_pe_ was found to remain largely invariant for a given species while being distinctly different from other species. This could be representative of distinct wall loosening mechanisms. For instance, cells of higher plant cells such as the pea stem, *P. sativus L.* (Π_pe_ = 32), loosen their walls by disrupting the hydrogen bonds between microfibrils using the protein, expansin ^40^. In the algal internode cells of *C. corallina* (Π_pe_ = 564), calcium bridges are broken and reformed between pectin polymers to loosen the wall ^32,41^, while the mechanism of loosening is still largely unknown for the sporangiophores of *P. blakesleeanus* (Π_pe_ = 1865) ^27,42^. However, the Π_pe_ values were found to be significantly different for these species by orders of magnitude. Furthermore, it has been shown that using a single approximately invariant value of Π_pe_ for a given species, one can determine variations in growth rate, *v_s_*, when the wall dynamics is altered by growth conditions. For example, different growth rates were correctly predicted for the cells of pea stems (i) “incised and growing in water”, and (ii) added with growth hormone IAAA, by using the same Π_pe_ value. The sporangiophores of *P. blakesleeanus* in growth stages I and IV show different growth rates but have approximately the same Π_pe_ values. Given the importance of Π_pe_, it is noteworthy that Eq. 15 provides a direct link to the molecular average bond detachment rate, k*_d_*, which can provide crucial insight into wall loosening mechanisms of different species given their molecular timescale. In the following section, we investigate the expansive growth behavior of three species of walled cells with distinct Π_pe_ values to estimate the varying molecular time scales corresponding to different wall loosening mechanisms.

In a recent review of mathematical models of expansive growth of cells with walls ^1^, it was recommended that all future global and local mathematical models should be able to reproduce the experimental results of *in vivo* stress relaxation and *in vivo* creep experiments in order to provide evidence that the model obeys the underlying physics and constitutive relationship measured experimentally. Thus, we investigate three types of mechanical behavior of the cell wall namely (a) steady growth or creep (irreversible deformation) where the wall force and creep velocities are constants, (b) stress relaxation when the wall length is held constant, and (c) a sudden step-up in wall force that produces a purely elastic (reversible) deformation followed by an increase in steady growth rate. These mechanical behaviors correspond to experiments conducted for plant, algal and fungal cells. Using the newly introduced statistical model of the cell wall, we now compare the model’s results with the experimental results from three distinct species of walled cells including *P. sativus* L. and *P. blakesleeanus*, and *C. corallina*. Using the equations derived for the global wall mechanics in section 2, the tether properties *k_d_*, *k_a_*, *K*, and *f_c_* are fitted to match experimental measurements. In the final part of this section, we present the mapping between molecular parameters of our statistical model and global wall mechanics that provide insights into regulation of the wall during growth.

### 3.1 Steady Growth (creep) and Stress Relaxation

During steady growth, the rate of irreversible wall deformation remains approximately constant in time if the turgor pressure does not change ^7^. This condition is discussed at the end of section 2.2 with the statistical model where a constant wall force, *F* = *PA*_0_, is applied to a population of tethers in the cell wall network. The solution to the continuum model for such a state is given in Eqs. 11 and 12, where a steady creep velocity, *v_g_*, can be estimated from 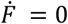.

The ability of the cell wall to loosen or relax is a crucial feature that allows for expansive growth. To quantify this behavior, let us now consider the cell wall growing at a rate, *v_g_*, to be suddenly arrested from further growth at time, *t* = *t*\*, i.e. *v*(*t* > *t*\*) = 0. The microfibrils stop moving while the detachment and attachment of tethers continue. Since tethers below the threshold force *f_c_*, cannot detach (Eq. 2), they remain intact with the stretch that was imposed at *t*\*. The rest of the attached tether population undergo the detachment process and re-attach in the stress-free state, *δ* = 0. The histogram in Fig. 4 from Monte Carlo simulations shows that at long times, all the tethers with elongation *δ* > *δ_c_* have detached and reattached at *δ* = 0. In the continuum limit, the distribution function *ϕ*(*δ*, *t*) is obtained from Eq. 11a by substituting *v* = 0 (see Appendix B.3 for detailed proof). For *t* > *t*\*, the distribution function in the region below the threshold elongation, *δ_c_*, remains unaffected from steady growth in Eq. 12, i.e. *ϕ*(*δ* < *δ_c_*, *t*) = *ϕ_s_*, while the other part decays exponentially to zero at rate *k_d_*. This process causes the total wall force *F*, to reduce with time, thus resulting in stress relaxation as seen from the Monte Carlo simulations and the continuum model in Fig. 4. By substituting *v*(*t* > *t*\*) = 0 in the continuum model (Eq. 11a), we obtain

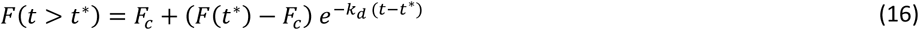

that predicts an exponential stress relaxation rate with a stress relaxation time, *t_R_* = 1/*k_d_*. The halftime of exponential decay, i.e. the time taken for the wall force to relax to half of its initial value, can be calculated from Eq. 16 as, *t*_1/2_ = ln 2 /*k_d_*. The half time is often useful in quantifying relaxation in experimental measurements and provides a time scale that can be compared for different species. It may further be noted that the wall force approaches the constant value *F_c_*, at long times (i.e. *t* → ∞) given in Eq. 11b as the sum of all tether forces below the threshold *f_c_* that remain attached.

**Figure 4:**
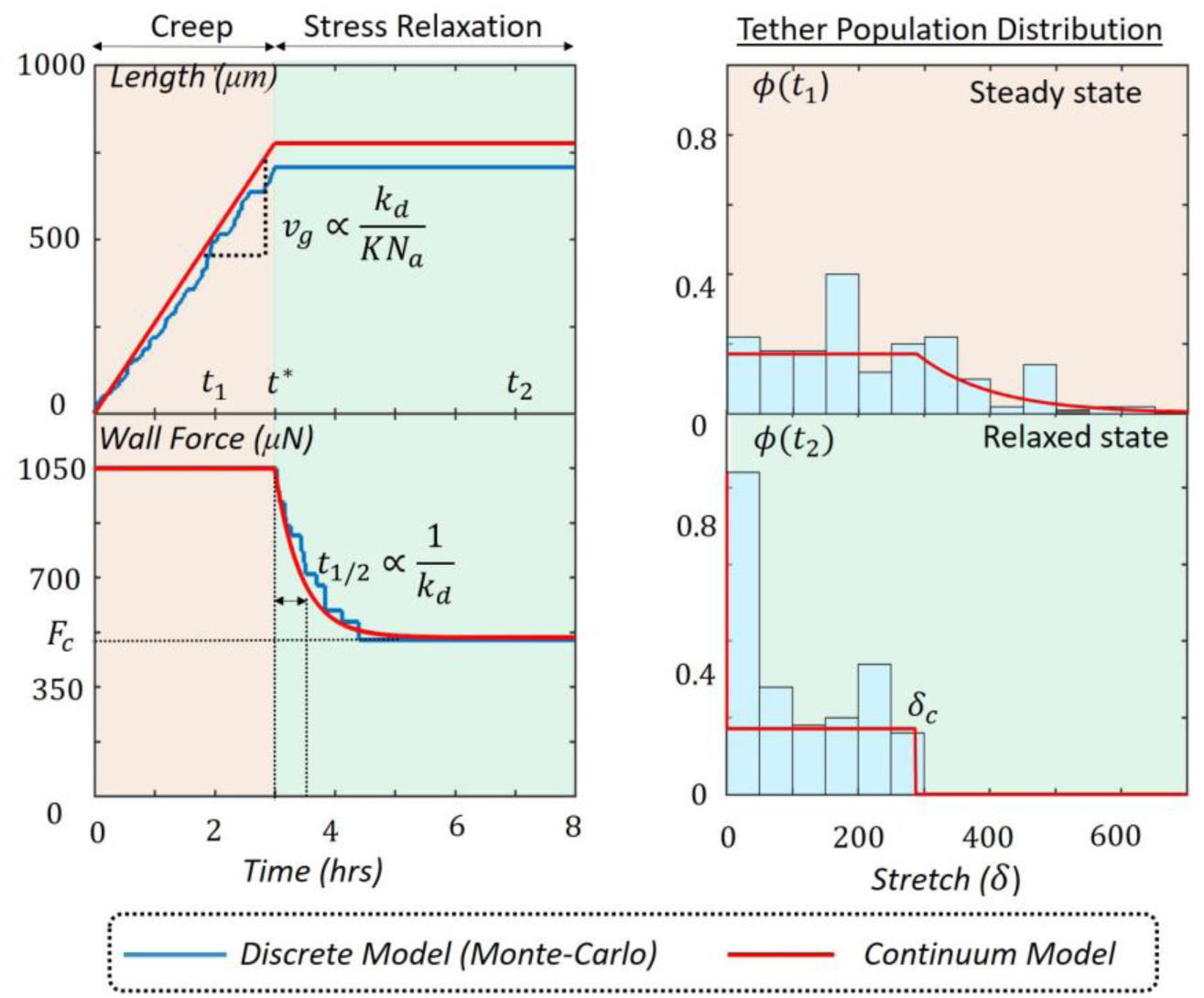
Illustration of the steady growth or creep (red panel) and stress relaxation (green panel) with Monte Carlo discrete simulations and the continuum model shown along with histograms and continuous distribution function ϕ. The population size for the Monte-Carlo simulations is N = 100. The molecular parameters used for all simulations are k_a_ = 10 h^−1^, k_d_ = 2 h^−1^, NK = 5 Nm^−1^ and Nf_c_ = 1500 μN.

### 3.2 Steady Growth and Step-up in Turgor Pressure

An alternative method for measuring the mechanical properties of the wall is through turgor pressure manipulation. These tests are typically performed on growing cells where the turgor pressure is increased suddenly (stepped-up) with a pressure probe. The steady growth rate on the other hand may be used to estimate the plastic or viscous properties of the cell wall (such as the extensibility *ϕ* of the Ortega equation, Eq. 13). Depending on the cell species, a more or less pronounced elastic response can be observed during pressure step-up; measuring this elastic deformation thus provides an interesting method by which one can estimate the elastic properties of the wall (the elastic modulus *ϵ* from the Ortega equation, Eq. 13). Let us therefore consider the condition of a sudden increase in turgor pressure during which the wall force is increased from *F* at time *t* = *t*\* to *F* + Δ*F* at time *t* = *t*\* + *dt*. The time period *dt* is here taken to be extremely short compared to the average lifetime 1/*k_d_* and so that tether detachment and attachment do not take place and the wall deformation remains purely elastic. Since the wall stiffness corresponds to the combined effective value of all attached tethers, *KN_a_*(*t*\*), at the time of the pressure step-up, the elastic elongation in wall length Δ*l* can be easily determined as:

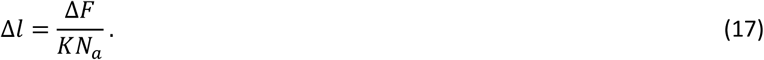

**Figure 5:**
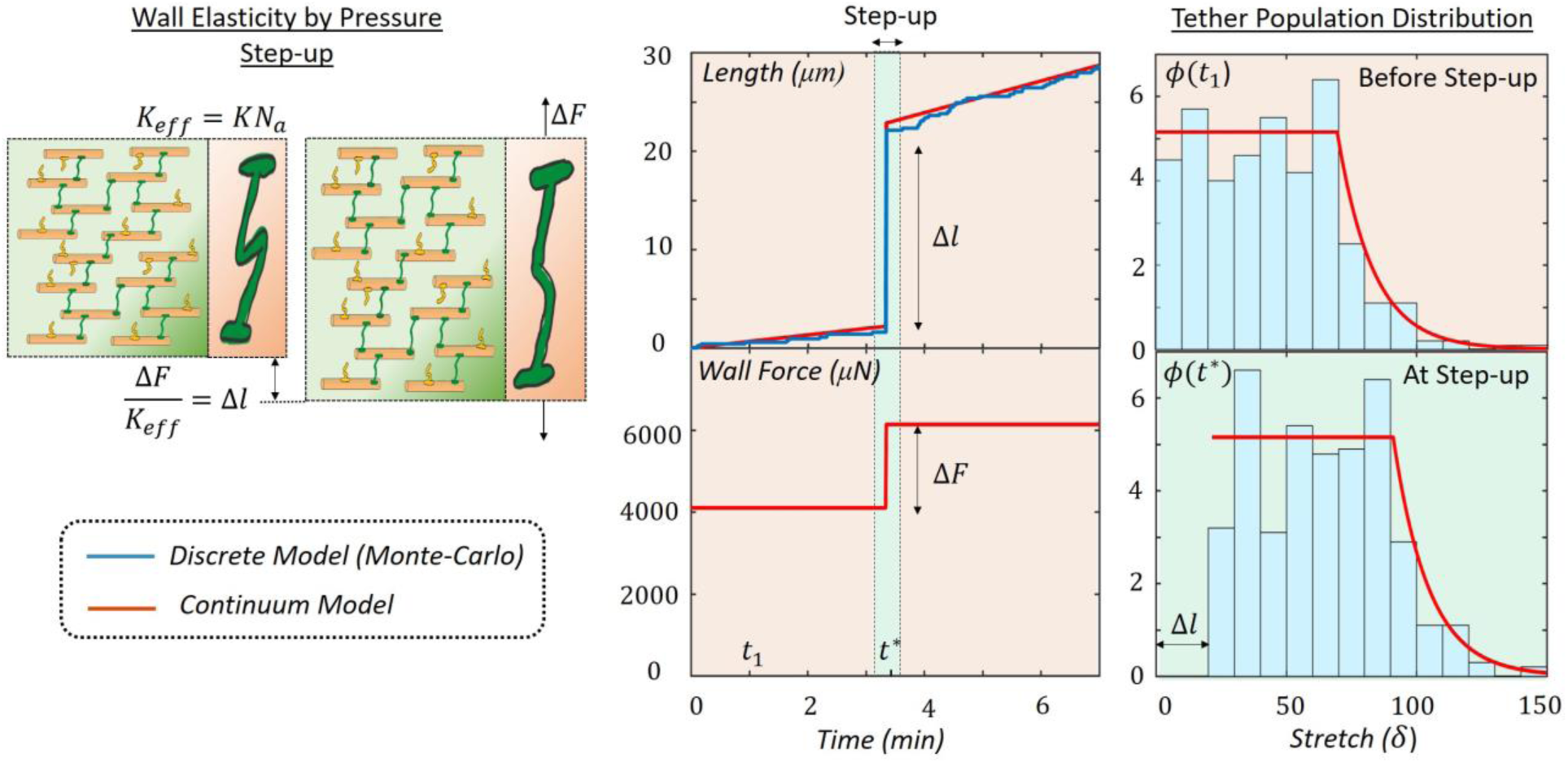
Left: Illustration of the network elasticity during a pressure step-up. Right: Illustration of the steady growth or creep (red panel) and step-up (green panel) with Monte Carlo discrete simulations and the continuum model shown along with histograms and continuous distribution function ϕ. The population size for the Monte-Carlo simulations is N = 500. The molecular parameters used for all simulations are k_a_ = 2.2 h^−1^, k_d_ = 3 h^−1^, NK = 115 Nm^−1^ and Nf_c_ = 8000 μN.

The elastic elongation Δ*l* is also the change in the length of each individual tether in the population that remains attached at *t* = *t*\*. This can be observed from the distribution function, *ϕ*(*t*\*), obtained from discrete Monte-Carlo simulations and the continuum model as show in Fig. 5 for a test case with given molecular parameters (see Fig. 5 caption). Compared to the distribution function at a time *t*_1_ < *t*\*, the step-up in wall force shifts the distribution function to the right by an amount equal to Δ*l*.

### 3.3 Comparison with Experimental Measurements

An advantage of the presented approach is that it enables to estimate molecular quantities, such as kinetics and mechanical properties based on macroscopic experiments. We here engage in such a study based on experimental measurements of the steady growth and stress relaxation in incised pea stem (*P. sativus L.)* ^13^ and the fungal cell, *P. blakesleeanus* in growth stage IVb ^7^. Stress relaxation experiments are conducted by removing the cell from its water supply and preventing its transpiration. The removal from water involves incision from the tissue or direct removal from the mycelium. The pressure decay is then measured with a pressure probe to determine the stress relaxation characteristics including the rate and final pressure value after decay *P_c_*. The turgor pressure decay for *P. sativus L.* and *P. blakesleeanus* are shown in Fig. 6 where the latter relaxes significantly faster than the former. The molecular parameter *k_d_* is determined from the measured half time, *t*_1/2_, while *k_a_*, *K* and *f_c_* are fitted with the growth rate before relaxation, *v_g_*, and the critical pressure, *P_c_*, with good agreement (Fig. 6). Since the wall areas of *P. sativus L.* (*R* = 23.5 μm and *l* = 3.5 mm) and *P. blakesleeanus* IVb (*R* = 75 μm and *l* = 30 mm) vary in orders of magnitude, tether stiffness and threshold force are normalized to the wall area. In other words, the fitted tether properties are reported as *nK* and *nf_c_* respectively (see Table 1), where *n* is the number of tethers per unit wall area. The fitted values of parameters, *k_a_*, *nK* and *nf_c_*, based on measurements of *v_g_* and *P_c_*, are found to be non-unique. For instance, the same value of *P_c_* can be reached by either considering many attached tethers (*N_a_*) with a small force threshold (*f_c_*) or considering fewer attached tethers with a high force threshold. This implies that a wide range of values of *k_a_* and *nf_c_* is possible as they determine the value of *N_a_* and *P_c_*. The values of threshold force, *nf_c_*, are significantly higher for cells of pea stems than the sporangiophores of *P. blakesleeanus* and which could explain the difference in the values of *P_c_*. Comparison of the tether stiffness, *nK*, remains inconclusive as a wide range of values are possible for *P. blakesleeanus* (Table 1). The values of Π*_pe_* can be calculated for both species based on the values of *k_d_* fitted with the above experiments and are found to be 33.7 and 1030 for pea and the sporangiophores respectively. These numbers are in good agreement with values reported in literature ^23^ which are 32 ± 6 and 1246. The *in vivo* stress relaxation test of steady growing cells, therefore, provides a method to determine certain molecular properties (such as *k_d_*) with good accuracy while other properties (such as *K* and *f_c_*) can only be estimated with low accuracy. We note that the estimation of the attachment rate *k_a_* has the highest uncertainty in fitting with this test.

**Figure 5:**
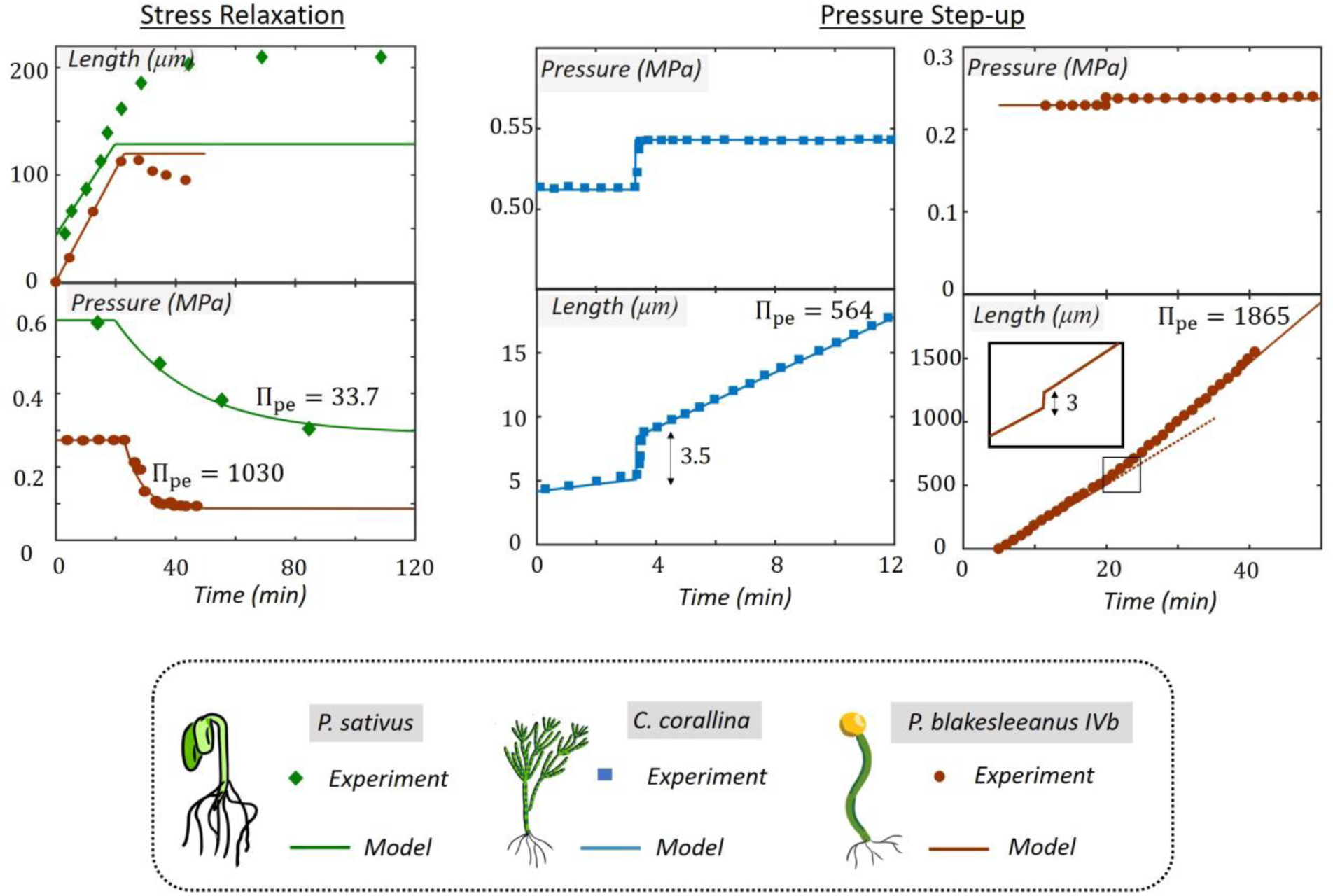
Left: Comparison of experimental measurements and model estimation of pressure relaxation of pea stem, P. sativus L. ^13^ and fungus, P. blakesleeanus ^7^. Note that the magnitudes of Π_pe_ and k_d_ are different for P. sativus L. and P. blakesleeanus for the stress relaxations. Right: Comparison of experimental measurements and model estimation of steady growth before and after pressure step-up in the algal cells C. corallina ^19^ and fungus, P. blakesleeanus IVb ^7^.. Note that the magnitudes of Π_pe_ and k_d_ are different for C. corallina and P. blakesleeanus.

**Table 1:**
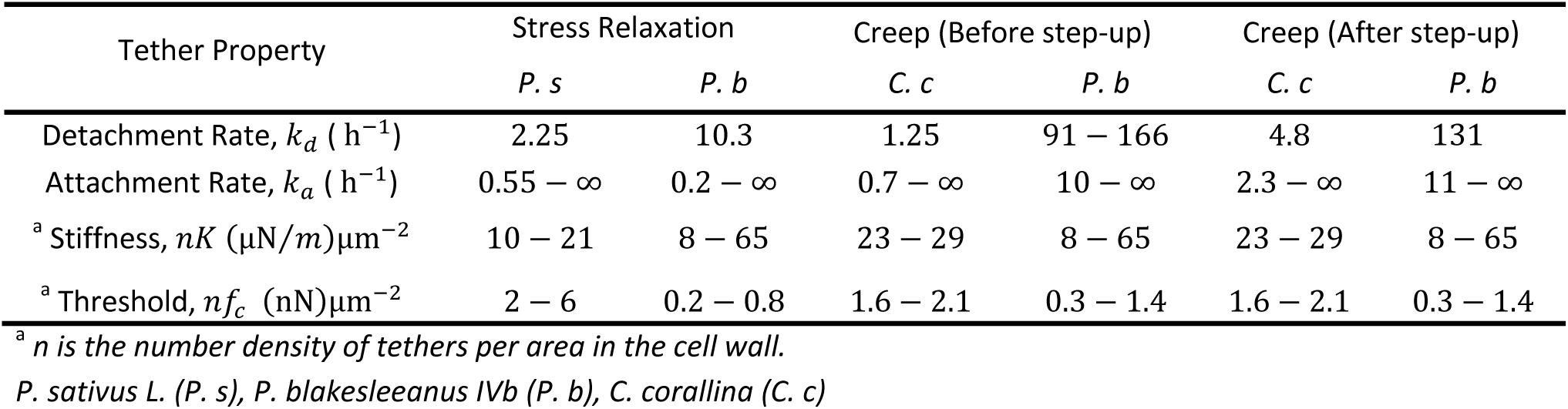
Fitted tether properties for model prediction of stress relaxation and step-up experiments

| Tether Property | Stress Relaxation |  | Creep (Before step-up) |  | Creep (After step-up) |  |
| --- | --- | --- | --- | --- | --- | --- |
|  | *P. s* | *P. b* | *C. c* | *P. b* | *C. c* | *P. b* |
| Detachment Rate, $k_d$ ($\text{h}^{-1}$) | 2.25 | 10.3 | 1.25 | 91 – 166 | 4.8 | 131 |
| Attachment Rate, $k_a$ ($\text{h}^{-1}$) | 0.55 – ∞ | 0.2 – ∞ | 0.7 – ∞ | 10 – ∞ | 2.3 – ∞ | 11 – ∞ |
| <sup>a</sup> Stiffness, $nK$ ($\mu\text{N}/\text{m}$)$\mu\text{m}^{-2}$ | 10 – 21 | 8 – 65 | 23 – 29 | 8 – 65 | 23 – 29 | 8 – 65 |
| <sup>a</sup> Threshold, $nf_c$ (nN)$\mu\text{m}^{-2}$ | 2 – 6 | 0.2 – 0.8 | 1.6 – 2.1 | 0.3 – 1.4 | 1.6 – 2.1 | 0.3 – 1.4 |

<sup>a</sup> $n$ is the number density of tethers per area in the cell wall.
*P. sativus* L. (*P. s*), *P. blakesleeanus* IVb (*P. b*), *C. corallina* (*C. c*)

Molecular parameters for the algal cells, *C. corallina* and the sporangiophores of *P. blakesleeanus* in stage IV (Table 1) are then estimated using both *in vivo* creep and pressure step-up response. For this, we proceed in two steps. First, we estimate *f_c_*, *k_d_*, and *k_a_* by matching the critical turgor pressure *P_c_*, available from other experiments ^7,19,20^ and the steady growth rate *v_g_*. Second, we estimate the tether stiffness *K* using the elastic response of *C. corallina* from Eq. 17. We note here that the elastic response due to step up is measurable in *C. corallina* while this is not the case with *P. blakesleeanus* (see Fig. 6) where the elastic deformation is negligible. However, based on independent measurements of the elastic modulus ^44^, *ε* = 60.9 ± 5.1 (SE)MPa, the range of possible values for tether stiffness, *K*, is shown in Table 1. As in the case of stress relaxation, we find that *in vivo* creep measurements also lead to non-unique fits for model parameters. Fortunately, with previously reported ^23^ values of Π*_pe_* = 1865 ± 583, it is possible to confine the range of values of *k_d_* using Eq. 15 to that reported in Table 1.

Despite the above difficulties, distinct differences can still be observed between the two cell types in the magnitudes of molecular parameters. First, we note that growth of the *C. corallina* cells is significantly slower than *P. blakesleeanus*; this is reflected in the estimated values for tether dynamics (*k_a_* and *k_d_*) that are higher by an order of magnitude in *P. blakesleeanus*. Second, the critical pressure *P_c_* is reported from previous experiments to be higher in *C. corallina* (*P_c_* =0.35 MPa ^20^) than *P. blakesleeanus* (*P_c_* = 0.085 MPa ^7^). Accordingly, the critical force for dissociation, *f_c_*, of tethers is found to be higher for *C. corallina*. Based on our fitted parameters, we show that the model prediction of the elastic response due to pressure step-up is significantly smaller (inset in Fig. 6) in *P. blakesleeanus*. Though the absolute value of the predicted elastic response (Δ*l* = 3 *μm*) is comparable to that measured for *C. corallina* (Δ*l* = 3.5 *μm*), the relative value to the length of the cell, *l*, is more significant in the latter than the former.

Finally, we note that the creep rate after pressure step-up in both cell types is estimated by the model with a higher value of *k_d_* (See Table 1). This estimate is obtained using Eq. 15 from which the steady growth rate is given by *v_g_*/*l* = Π_pe_*k_d_*, where Π_pe_ is approximately constant for a given species irrespective of growth rate ^23^. The higher estimate of *k_d_* could indicate the possibility of a nonlinear relationship between wall force and creep rate that is not currently addressed in our model. It is likely that the tether dissociation rate, *k_d_*, is amplified by force making the creep rate higher when the wall stress is increased due to turgor pressure. This may be investigated in future work with the proposed statistical model where *k_d_* in Eq. 2 may be modified to include a force-dependence, and more specifically, an increase in dissociation rate with tether force.

### 3.4 Tuning Tether Dynamics to Regulate Wall Mechanics

Having compared the model’s extension and turgor pressure behavior against relevant experimental results and obtaining good agreement, we now propose to examine how the proposed statistical approach may help better understand the effect of the molecular scale on growth dynamics. We particularly focus on four key parameters, namely *k_d_*, *k_a_*, *K*, and *f_c_*.

For this, let us consider the steady growth (at rate *v_g_*) of a cell wall under a constant turgor pressure *P*. From Eq. 11a, the contributions of tether stiffness, *K* and detachment rate, *k_d_*, to the growth rate are apparent. However, the dependence of *N_a_* and *F_c_* on the other tether properties is not straightforward. Based on a parametric analysis of the solutions to the continuum model from Eqs. 11-12 (see Appendix B.2), we find that the ratio of attached tethers in the total population, *N_a_*/*N*, is inversely correlated to the ratio, *k_d_*/*k_a_* (Fig. 7a). In other words, most tethers are attached to the fibrils at steady state when *k_a_* is more dominant than *k_d_* and vice-versa. Interestingly, the wall yield force *F_c_* is also inversely correlated to *k_d_*/*k_a_*. This implies that for a given wall force or turgor pressure, altering the tether dynamics, i.e. raising or lowering *k_d_*/*k_a_*, has a compound effect on the growth rate, *v_g_*, by affecting both *N_a_* and *F_c_*. The distribution function for steady growth, *ϕ*(*δ*, *t*), further shows that when the ratio *k_d_*/*k_a_* is high, both *N_a_* and *N_s_* (number of tethers below the critical force) are low. This explains the reduction in wall yield stress, *F_c_* at large values of *k_d_*/*k_a_* though the threshold force *f_c_* remains unchanged. Alternately, if the critical force of each tether, *f_c_*, is increased, it is predicted that, at steady state, there exists a high fraction of attached tethers and hence a high wall yield force, *F_c_* (Fig. 7a). It is worth noting that from the plots in Fig. 7a, each combination of values of *N_a_* and *F_c_* can be obtained by many different values of *k_d_*/*k_a_* and *f_c_*. This explains the problem of non-unique fitting parameters noted in the previous sections for experimental measurements of steady growth.

**Figure 7:**
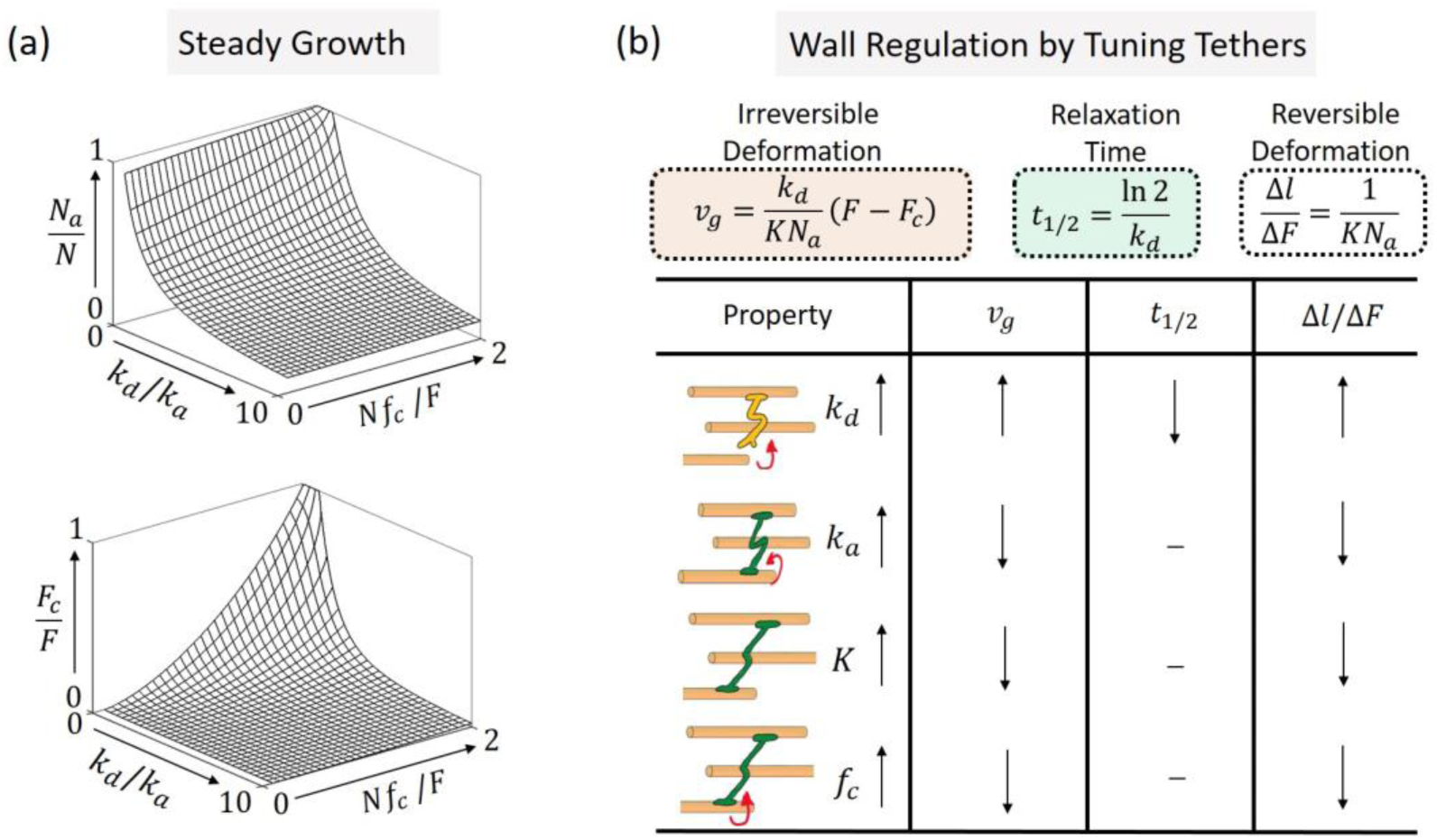
(a) The relationship between the ratio of tether detachment and attachment rates, k_d_ and k_a_, and tether threshold force, f_c_ with the fraction of tethers, N_a_/N, and the fraction of wall yield force, F_c_/F, and (b) Summary of the roles of the four tether properties, k_d_, k_a_, K and f_c_ on the growth rate, relaxation half time and the reversible deformations.

The roles of the four tether properties in the steady growth rate or irreversible deformation are thus summarized qualitatively in Fig. 7b. While increasing *k_d_* causes faster growth, increasing the other three parameters hinders growth by slowing down the wall deformation rate for a given turgor pressure. The effect on reversible deformations, (e.g. during a turgor pressure step-up experiment) follows the same trend as irreversible deformations. This indicates that for a given wall force, the irreversible deformation or growth rate cannot be regulated to be fast without making the wall “soft” (either soft or few attached tethers) and vice-versa. The half time of relaxation, on the other hand, is governed only by the detachment rate, *k_d_*, and is independent of all other parameters, which do not play a role in dissipating energy. Importantly, *k_d_* can be determined for each walled cell if the magnitudes of Π*_pe_* and *v_s_* are known for that cell (see Eq. 15).

## 4 DISCUSSION AND FUTURE RESEARCH

Here, we developed a local numerical-mathematical model that explicitly employs the same mechanism used by cell walls to manipulate their mechanical properties, i.e. by breaking load-bearing (stress-bearing) molecular bonds and remaking bonds without stress (zero load). More specifically, we introduced a probabilistic model to describe the dynamics of the tethered network of microfibrils in the cell wall during expansive growth. In this framework, the irreversible wall deformation is explained by the intermittent detachment and re-attachment of elastic tethers, while reversible deformations originate from their elastic elongation. A statistical mechanics was then used to describe the time evolution of the stretch distribution of a population of tethers that could then be used to estimate the evolution in wall forces. Based on this approach, a statistically based growth equation was derived, which provides a molecular interpretation of the established Ortega equation that describes the expansive growth of the cell wall. The extension rate and turgor pressure behavior produced by the statistical model was compared to experimental results from *in vivo* stress relaxation and creep with pressure steps of three different cell types, namely pea stem (*P. sativus L*.), algal internode cells (*C. corallina*) and sporangiophores of *P. blakesleeanus,* with good agreement.

It is shown that the statistical model’s extension behavior as a function of applied force is dependent predominately on the magnitude and behavior of four variables; *k_d_*, *k_a_*, *K*, and *f_c_*. Modeling the force (turgor pressure) and extension (growth) behavior for any walled cell requires fitting the magnitude (and behavior) of these four variables. A section is devoted to explaining how fitting these variables to produce the desired force-extension behavior are non-unique and guidelines are provided. In general, it is found that the magnitude of two of the variables (*k_d_* and *f_c_*) can be obtained from the results of *in vivo* stress relaxation experiments and the magnitude of *K* can be estimated from the magnitude of the volumetric elastic modulus. An important finding is that the dimensionless number Π_pe_ is related to *k_d_* and *v*_s_: Π_pe_ = (*k_d_*/*v_s_*) (Eq. 15). This relationship allows us to calculate the magnitude of *k_d_* when Π_pe_ and *v*_s_ are known. A good fit is obtained for the turgor pressure step-up experiments by maintaining a constant value for Π_pe_ and thus changing *k_d_* when *v*_s_ changes. The principle of keeping Π_pe_ constant for a single cell is consistent with the experimental findings that Π_pe_ is approximately constant for any one species of walled cell but different for other species ^23^.

The proposed statistical model has the potential to unlock avenues of research that relate the magnitudes of *k_d_* and *k_a_* with specific enzymes and proteins that are found to loosen walls in different species of walled cells. If similar research identifies the magnitudes of *K* for tethers used in cell walls, and bond strength, *f_c_*, for load bearing bonds, then it is envisioned that a statistical model using this platform can be used to conduct parametric studies to identify which combination of enzymes, proteins, tethers, and molecular bonds are used to obtain the force-extension behavior observed *in vivo*. Accurate estimation of these molecular properties will enable a better understanding of the wall regulation mechanisms that cells use under different growth conditions and stimuli responses to light, gravity, stretch, etc. ^45–47^ Furthermore, the framework of this model is powerful as it is based on simple rules to describe the dynamics of tether behavior and when applied to a population of tethers, it can predict the growth phenotype of different cells. Thus, the model offers ripe ground for the investigation and discovery of new mechanisms in cell morphogenesis during growth, such as helical growth in *P. blakesleeanus* ^35,48^ and apical tip growth seen in pollen tubes, root hairs, and algal rhizoids^2,49–51^.

## AUTHOR CONTRIBUTIONS

FJV designed the theoretical aspect of the study with the help of JKO who provided insights from experimental and theoretical expertise. SLS and FJV developed the statistical model and equations. SLS performed all model simulations and fits. All authors contributed equally to writing the manuscript.

## ACKNOWLEDGMENTS

FJV acknowledges the support of the National Science Foundation under CAREER award 1350090.

## APPENDIX A

### A.1 Methodology for Monte-Carlo Simulations

An assembly of springs are modelled to undergo deformation, attachment and detachment based on probabilities defined in Eq. 1. As the attachment and detachment events are assumed to follow a Poisson process ^52^, a better way to describe them is through the probability that a state (attached or detached) changes in a time interval *dt*. Therefore, the probability that an attached tether becomes detached in time *dt* is given by *k_d_dt*. Similarly, the probability that a detached tether becomes reattached in time *dt* is given by *k_a_dt*. The algorithm for the simulating the dynamics of the tether network by the discrete approach (Monte-Carlo) is summarized in the steps below.

*<u>Input</u>:* Number of tethers {*N*}, Molecular parameters {*k_a_*, *k_d_*, *K*, *f_c_*}

*<u>Output:</u>* Deformation speed {*v*(*t*)} (creep), Wall force {*F*(*t*)} (stress relaxation)

#### Algorithm

**1.** Initialize tether state, *s_i_* (0 when detached and 1 when attached), and tether elongation *δ_i_* = *F*(0)/∑(*Ks_i_*), *i* = 1: *N*.
**2.** Start time loop with *τ* = 1 to *τ_max_* with time step *dt*

**2.1** Start attached tether population loop, *j* = 1: *N_a_*

- Sample a random number, *r_j_*, between 0 and 1.
- if *r_j_* < *k_d_dt*

- Assign *s_j_* = 0
- Store force lost due to detachment, 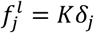 End attached tether population loop
**2.2** Start detached tether population loop, *k* = 1: *N* − *N_a_*

- Sample a random number, *r_k_*, between 0 and 1.

- if *r_k_* < *k_a_dt*

- Assign *s_k_* = 1
- Assign *δ_k_* = 0 End detached tether population loop
**2.3** Compute updated force from the network, *F*(*τ*) = ∑*_i_*_=1:*N*_*Kδ_i_*(*τ*)
**2.4** Compute change in deformation of the wall, 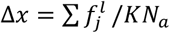
**2.5** Update overall deformation of the wall, *x*(*τ*) = *x*(*τ* − 1) + Δ*x*
**3**. End time loop

## APPENDIX B

### B.1 Derivation of the Macroscopic Force-Velocity Equation

The relationship between velocity and force in Eq. 11a is derived from the integral equation for force given in Eq. 5b by differentiating w.r.t time as

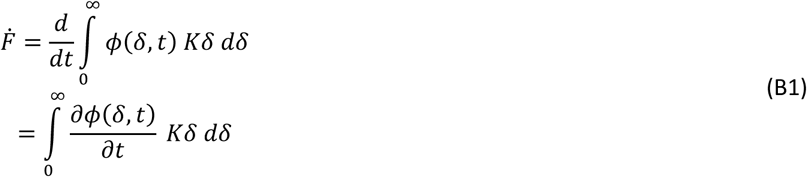

Using the evolution equation Eq. 10, we can re-write the rate of change of force as

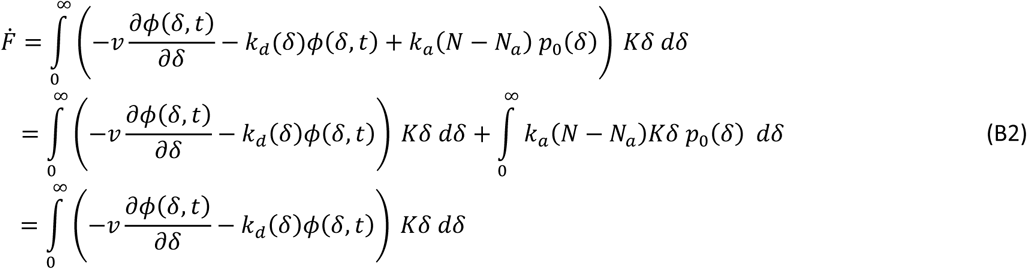

where using the property of Dirac delta function, *p*_0_(*δ*), the integral of the attachment term vanishes. Further, using integration by parts for the first term in the integrand, we get

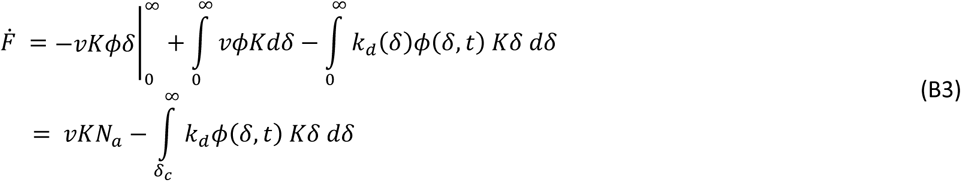

where Eq. 5a is used for the first integral term and Eq. 2 is used for the second integral term to simplify it into two different domains namely, (0, *δ_c_*) and (*δ_c_*, ∞), such that *k_d_*(*δ*) = 0 in the first domain. The boundary term vanishes as a result of the condition that lim_δ→∞_ *ϕ*(*δ*)*δ* = 0, assuming an Gaussian decay of *ϕ*(*δ*) to zero at large *δ* values. Rewriting the second term of Eq. B3 using Eq. 5b, we obtain

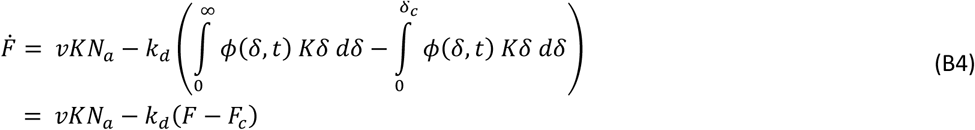

where the critical wall force, *F_c_*, is given in Eq. 11b.

### B.2 Derivation of the Steady State Creep Solution

Let us consider evolution equation of the tether stretch distribution, Eq. 10. The last term, *k_a_*(*N* − *N_a_*) *p*_0_(*δ*), that corresponds to the tether re-attachment always contributes to the distribution function at *δ* = 0, as *p*_0_(*δ*) is assumed to be a Dirac delta function. Therefore, this term can be re-formulated as a source boundary condition at *δ* = 0 for *ϕ*. Let the number of attached tethers at the stress-free configuration, *δ* = 0, be denoted by *N*_0_. The time rate of change of *N*_0_ is governed solely by attachment events and therefore can be written as

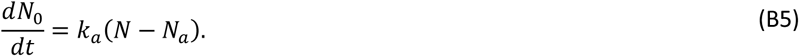

Now, using *dN*_0_ = *ϕ*(0, *t*) *dδ*, and the definition of constant velocity of microfibrils, *v_g_* = *dδ*/*dt*, the above equation gives

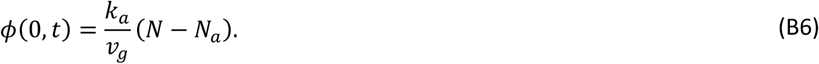

With the above boundary condition at *δ* = 0, the evolution equation, Eq. 10 reduces to

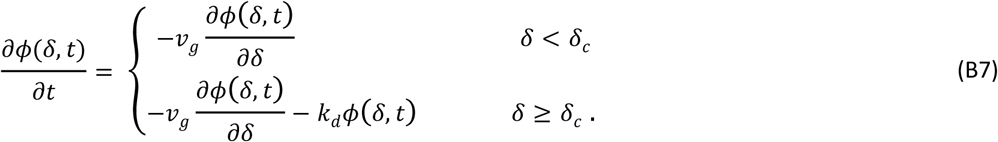

At steady state creep, *∂ϕ*/*∂t* = 0. Using Eqs. B6-B7, *ϕ*(*δ*, *t*) can then be found by solving the PDE analytically in each domain, *δ* < *δ_c_* and *δ* ≥ *δ_c_*, with the condition that the function *ϕ*(*δ*, *t*) is continuous at *δ* = *δ_c_*. The solution is shown in Eq. 12.

Using the solution in Eq. 12, the number of tethers below the threshold elongation (*δ_c_*), denoted by *N_c_*, and the critical wall force, *F_c_*, are now given by

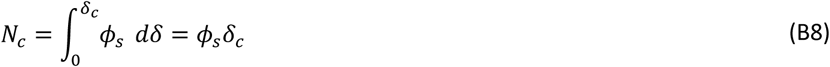

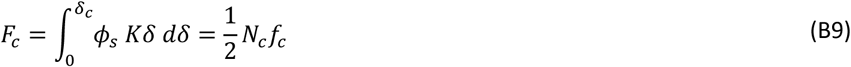

where *ϕ_s_* = *k_a_*(*N* − *N_a_*)/*v_g_*. The number *N_c_* can also be obtained by integrating the evolution equation, Eq. 10, across all values of *δ* from 0 to ∞, to give

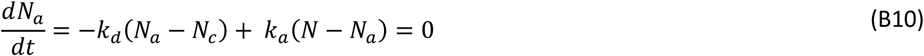

at steady-state creep.

Therefore, solving the algebraic equations in Eq. B8-B10 along with Eq. 11a with 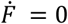, gives the following expression for the fraction of attached tethers, *N_a_*/*N*, and critical wall force, *F_c_*, as

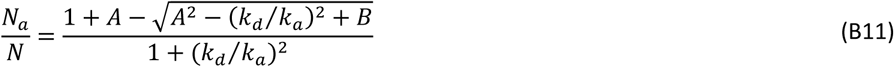

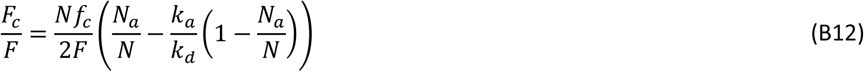

where the terms *A* = (*F*/*Nf_c_*)(*k_d_*/*k_a_* + *k_d_*/*k_a_*)^2^), and *B* = (2*F*/*Nf_c_*)((*k_d_*/*k_a_*)^2^ − (*k_d_*/*k_a_*)^3^).

### B.3 Derivation of the Distribution Function during Stress Relaxation

When a network undergoing steady creep at velocity, *v_g_*, is suddenly stopped from further elongation at time, *t* = *t*\*, the evolution equation of the distribution function Eq. B7 is modified for time *t* > *t*\* as

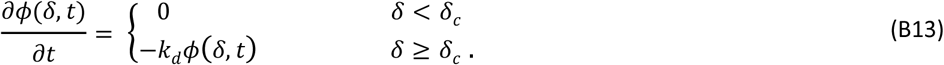

where the initial condition is given by

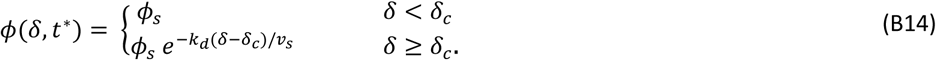

Eq. B13 shows that the distribution function does not change with time for tethers with elongation *δ* < *δ_c_*. The above equations B13-B14 are solved analytically to give

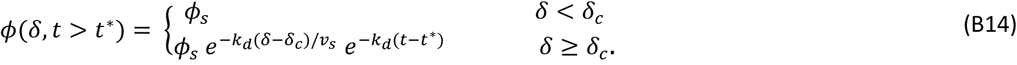

that describes an exponential decay of the distribution function with rate *k_d_*.

